# Unlocking the power of gene banks: diversity in base growth temperature provides opportunities for climate-smart agriculture

**DOI:** 10.1101/2024.11.07.622441

**Authors:** Clara Gambart, Jelle Van Wesemael, Rony Swennen, François Tardieu, Sebastien Carpentier

## Abstract

Implementation of context-specific solutions, including cultivation of varieties adapted to current and future climatic conditions, were found to be effective in establishing resilient, climate-smart agricultural systems. Gene banks play a pivotal role in this. However, a large fraction of the collections remains neither genotyped nor phenotyped. Hypothesising that significant genotypic diversity in *Musa* temperature responses exists, this study aimed to assess the diversity in the world’s largest banana gene bank in terms of base temperature (T_base_) and to evaluate its impact on plant performance in the East African highlands during a projected climate scenario. 116 gene bank accessions were evaluated in the BananaTainer, a tailor-made high throughput phenotyping installation. Plant growth was quantified in response to temperature and genotype-specific T_base_ were modelled. Growth response of two genotypes was validated under greenhouse conditions, and gas exchange capacity measurements were made. The model revealed genotype-specific T_base_, with 30 % of the accessions showing a T_base_ below the reference of 14 °C. The Mutika/Lujugira subgroup, endemic to the East African highlands, appeared to display a low T_base_, although within subgroup diversity was revealed. Greenhouse validation further showed low T sensitivity/tolerance to be related to the photosynthetic capacity. This study, therefore, significantly advances the debate of within species diversity in temperature growth responses, while at the same time unlocking the power of gene banks. Moreover, we provide a high throughput method to reveal the existing genotypic diversity in temperature responses, paving the way for future research to establish climate-smart varieties.

## 1 Introduction

Agriculture is more than ever challenged to increase food, feed, fuel and fibre production, without degrading the environment or impeding the future production potential (Foley et al., 2011). Climate-smart agriculture encourages the implementation of flexible, context-specific solutions that increase the resilience and resource use efficiency of agricultural systems (Lipper et al., 2014). Increased crop and variety diversity, along with improved pest, water and nutrient management have been proposed as valuable opportunities to maintain yield and alleviate yield gaps (Anderson et al., 2020; Zorrilla-Fontanesi et al., 2020).

The 11 international gene banks of the Consortium of International Agricultural Research Centres (CGIAR), conserving germplasm collections of 22 mandate crops, can play a pivotal role in the climate-smart agriculture framework by recommending accessions for breeding or by distributing and introducing the varieties that possess advantages for the local conditions. However, a large fraction of the managed germplasm is neither genotyped nor phenotyped (Van den houwe et al., 2020).

One of the conserved mandate crops is banana (*Musa* spp.). It is a tropical C3 crop, currently cultivated in a wide range of climates in tropical and subtropical regions of Asia (43.2 % of the global production), Africa (32.5 %) and America (22.9 %) (FAO, 2023; Varma and Bebber, 2019). They are consumed far beyond their cultivation area. With an annual export of 26.4 million tonnes, banana was the most important tropical export fruit in terms of quantity and monetary value in 2021 (FAO, 2023). However, 84.5 % of the global annual production (equalling 143.6 million tonnes) is consumed locally (FAO, 2023). The East African highlands, a densely populated area in the African Great Lakes region, is the largest banana consuming region in the world (Karamura et al., 1998). The region is characterized by a hilly landscape, ranging from 900 to 2,000 meters above sea level (m.a.s.l.), with peaks up to 5,800 m.a.s.l. (WorldClim, 2022). Average temperature is relatively low compared to other tropical areas, ranging from 17 to 25 °C, and decreases with increased elevation (WorldClim, 2022). With a generally accepted reference base temperature (T_base_) of 14 °C (Ganry, 1980; Ganry and Meyer, 1975), only few varieties are currently known to thrive relatively well above 1,500 m.a.s.l. (Kamira et al., 2016; Ocimati et al., 2014). Suboptimal mean temperatures result in higher sucker development, increased crop cycle duration, reduced bunch weights and therefore, serious yield reductions (Kamira et al., 2016; Sikyolo et al., 2013; Turner et al., 2016).

Growing degree days (GDD) calculations have been extensively used in agriculture to predict plant phenology stages (Baraibar et al., 2018; Neild and Seeley, 1977). However, most studies use a crop-specific T_base_, explaining genotype-specific developmental rates by varying GDD rather than also taking into account a different T_base_. This underestimates the crop genetic potential in regions at higher elevation.

As host of the world’s largest banana (*Musa* spp.) collection, we hypothesise that significant genotypic diversity in temperature response exists that will enable future research to establish climate-smart varieties. Hence, this study aimed to (i) assess the *Musa* genotypic diversity in terms of T_base_ and (ii) investigate the impact of a projected climate scenario on the distribution and plant performance in the East African highlands. To do so, we designed the BananaTainer, a high throughput, tailor-made banana germplasm phenotyping installation (Carpentier and Eyland, 2020; Van den houwe et al., 2020)In contrast to field studies, which are time consuming and obscured by varying environmental conditions, our approach enables a fast pre-field screening of genotypic responses in a controlled environment (Negin and Moshelion, 2017). We present growth responses of different plant organs in function of temperature, models for the genotypic T_base_ of 116 banana genotypes and a validation of two contrasting growth responses under fluctuating greenhouse conditions.

## 2 Material and Methods

### 2.1 High throughput phenotyping in the BananaTainer

#### 2.1.1 Plant material and growth conditions

*In vitro* plantlets were obtained from the International Transit Centre (ITC) of the Alliance of Bioversity International and CIAT, hosted at KU Leuven (BE) and acclimatised to autotrophic conditions for 5 weeks in a growth room in hydroponics. Per run, 504 selected plants (30 plants per genotype, 42 plants per reference genotype) were transferred to the BananaTainer. In total more than 6,800 individual plants were grown and analysed.

The BananaTainer is a container (12 m x 2.4 m x 2.9 m) based, tailor-made, growth chamber (Urban Crop Solutions, Waregem, BE). The three level design allowed to grow plantlets up to a height of 55 cm (Supplementary Fig. S1). Plants were positioned according to a restricted randomised design, ensuring 10 to 14 plants per genotype per level. They were grown per six in an 80 cm x 60 cm tray holding 4 L of nutrient solution, adjusted from Swennen *et al*. (1986). Every 12 minutes the solution was refreshed with complete drain reuse. Filtering was ensured by a UV and 80 µm particle filter. The environment was set at 70 % relative humidity (RH), 400 ppm CO_2_ and 12 hour/12 hour (day/night) light regime. LED lights (Urban Crop Solutions, Waregem, BE) provided a spectrum relevant for banana (50:45:5 blue:red:far red) (van Wesemael et al., 2020). Plant growth was evaluated at several temperature (T) regimes: 3 level regimes programmed around an average T of 20 °C (25/15 °C 12 hour/12 hour day/night regime, vapour pressure deficit (VPD) of 0.45 kPa), 3 around 25 °C (30/20 °C, 0.55 kPa) and 3 around 30 °C (35/25 °C, 0.61 kPa) for 6, 5 and 4 weeks, respectively. Ambient temperatures and relative humidity were recorded in the BananaTainer by six loggers, two at each level (TESTO 175-H1, DE), resulting in an enhanced resolution of the screened T range between 18 and 30 °C (Supplementary Fig. S2). The regimes around 20 °C correspond to the climatic conditions during the rainy season in the East African highlands, the regimes around 25 °C mimic warmer conditions and the regimes around 30 °C represent predicted optimal conditions, based on the areas currently under dessert banana cultivation (Varma and Bebber, 2019). 116 accessions, belonging to 30 different subgroups were evaluated around 20 °C, 22 accessions around 30 °C of which 4 additionally around 25 °C (Supplementary Table S1). This subset covers a large genetic diversity, with special attention to the Mutika/Lujugira subgroup. Cachaco (ITC0643, ABB, Bluggoe subgroup) and Mbwazirume (ITC1356, AAA, Mutika/Lujugira subgroup) were chosen as reference genotypes, given their contrasting growth and stomatal responses to abiotic factors (Eyland et al., 2021; Kissel et al., 2015). They were present in each BananaTainer experiment.

#### 2.1.2 Growth quantification

Plant size was measured non-destructively at the start of every BananaTainer experiment. These measurements included canopy area, plant fresh weight, and height of the pseudostem. At the end of each experiment, plant growth was quantified both destructively and non-destructively. While non-destructive measurements were identical to the measurements at the start, destructive measurements consisted of fresh organ mass (leaves, pseudostem with rhizome and roots) and leaf area of the individual leaves.

Canopy and individual leaf areas were determined from topview RGB images with blue background and red reference surface of 50 cm^2^ (Supplementary Fig. S3). These pictures were analyzed, using a colour-based foreground-background segmentation based on SLIC superpixels (EBImage R-package, Olés et al. (2023)). As such, the green plant parts and the red reference surface of known area were segmented out of the image enabling leaf area extraction (Supplementary Fig. S3) (Eq. 1).

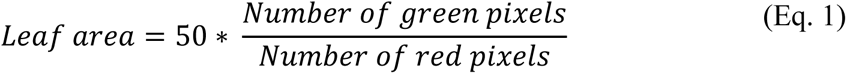

Given the different durations of the experiments, interpolation was used to estimate the morphological variables, related to plant development, plant growth and mass distribution at 4 weeks and compare the effects of the different treatments. As temperatures were kept stable during each experiment, leaves developed at a constant rate (Turner and Lahav, 1983). Moreover, plant mass increases proportionally with leaf area, enabling the estimation of all morphological parameters at 4 weeks. This resulted in a dataset with a total of 36 variables (crop ontologies as defined by van Wesemael et al. (2019); Supplementary Table S2).

#### 2.1.3 Base temperature (T_base_) estimation

To evaluate T_base_, genotype-specific canopy growth curves were established using the beta distribution model (Eq.2, Supplementary Fig. S4). This model has been successfully used to model yield (Ruiz-Vera et al., 2018; Varma and Bebber, 2019), vegetative (Devi and Reddy, 2018; Turner and Lahav, 1983) and photosynthetic responses (Adams et al., 2017) to temperature.

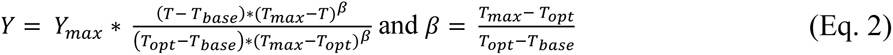

Y represents the weekly canopy growth, Y_max_ the maximum weekly canopy growth, T the temperature, T_base_, T_opt_ and T_max_ the base, optimal, and maximum temperature at which canopy growth occurs, respectively.

To reduce the variability in growth response, per level, every plant was assigned to the nearest temperature and relative humidity logger (Section 2.1) and the average genotypic growth response per logger was calculated. In this way six datapoints per experiment and genotype were obtained. If genotypes were screened in multiple experiments all datapoints were merged for growth modelling. Hence, growth models were based on 6, 12 and 18 datapoints for genotypes present in one, two and three experiments, respectively. Model parameters were estimated by minimizing the root mean square error (RMSE) based on the Fletcher variable metric method across 100 distinct initial values (Rvmmin R-package, Nash (2021)). Among the 100 models per genotype, only models below the accuracy threshold were retained. This threshold was determined by a Partial Least Squares Discriminant Analysis and decision tree modelling (Supplementary Fig. S5) (mixOmics R-package, Cao *et al*. (2021); tree R-package, Ripley (2022)). The genotype-specific model with minimum RMSE was selected as final growth model.

T_base_ precision is represented by the average standard deviation of the modelled T_base_ of the reference genotypes (Section 3.2). As these genotypes were present in each BananaTainer experiment, multiple growth curves were developed for all experiment combinations.

#### 2.1.4 Current and future climate impact predictions

The spatial impact of current and future climates on plant performance in the current banana growing region of the East African highlands was evaluated in two different ways, using qGIS (v3.16.1-Hannover; qGIS Development Team (2018)). First, raster calculations were made to visualize the number of genotypes with T_base_ larger than the average yearly T. Secondly, GDD raster calculations were made for two selected Mutika/Lujugira genotypes with contrasting T_base_, Guineo (ITC0005; AAA) and N’Dundu (ITC0732; AAA), to evaluate the annual heat unit accumulation above their genotype-specific T_base_ (Eq. 3):

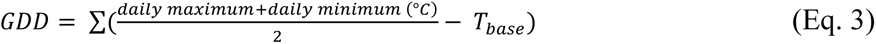

Global monthly current and future climate data was downloaded from WorldClim (2022) with a spatial resolution of 30 seconds (0.86 km^2^ at the equator). Future climate is based on the CMIP6 future climate projections with emission scenario SSP5-8.5.

### 2.2 Growth response validation in the greenhouse

#### 2.2.1 Plant material, growth conditions and quantification

Two Mutika/Lujugira genotypes with contrasting T_base_, i.e. Guineo and N’Dundu, were selected for further validation. Both genotypes were evaluated during two greenhouse experiments: one in December 2020 and one in March 2021 (average T of 17.1 °C and 25.3 °C, respectively). Frequency histograms of the experienced conditions are presented in Supplementary Fig. S6.

*In vitro* plantlets, obtained from ITC, were acclimatised to autotrophic conditions for 8 weeks in the greenhouse in 10 L pots containing peat-based compost. The plants were then randomly positioned on high precision balances (max weight of 12 kg ± 1 g) of the Phenospex platform (Phenospex, Heerlen, NL). This multi-lysimeter setup registers the systems’ weight (Eq. 4) every minute and couples it to a balance-specific watering unit, ensuring on a daily basis plant-specific irrigation (5 ml accuracy) until a water content of 2.3 g g^-1^ (1.83 pF). To avoid evaporation, the soil was covered by plastic. Therefore, the systems’ weight loss was only related to plant transpiration.

To follow up growth, pseudostem height was measured and topview images were taken weekly (Section 2.1.2) to evaluate canopy area (LA_topview_). At the end of every experiment fresh weight of the leaves, pseudostem with rhizome and roots were measured.

#### 2.2.2 Light- and CO_2_ response curves

Light- and CO_2_ response curves were developed for both genotypes (3 replicates per genotype) on the second fully expanded leaf using a LI-6800 with Multiphase Flash Fluorometer head (LI-COR, Nebraska, USA). Measurements were made in the morning to avoid afternoon stomatal closure (Eyland et al., 2021; van Wesemael et al., 2019).

Light response curves were developed after exposing the plants to 1500 µmol m^-2^ s^-1^ until a stable photosynthetic rate (A) was recorded. Afterwards PAR dropped to 0 µmol m^-2^ s^-1^ in several steps (1500 – 1200 – 900 – 600 – 500 – 450 – 400 – 350 – 300 – 250 – 200 – 150 – 50 – 0 µmol m^-2^ s^-1^) and a stable A was recorded at every level. The hyperbola-based and Ye function were applied to model light response for each plant using the excel routines of Lobo et al. (2013). Models with the lowest RMSE were chosen.

CO_2_ response curves were developed after exposure to 400 ppm CO_2_ and 750 µmol m^-2^ s^-1^, as this PAR was considered saturating by all light response curves. After stabilization of A, CO_2_ dropped to 30 ppm and afterwards increased to 1000 ppm in several steps (400 – 300 – 200 – 100 – 30 – 400 – 600 – 800 – 1000 – 400 ppm). CO_2_ response models were developed using the Excel routines of Sharkey and colleagues (Sharkey, 2016; Sharkey et al., 2007) to estimate the maximum carboxylation rate (V_cmax_), maximum electron transfer rate (J_max_) and maximum triose phosphate usage (TPU_max_).

### 2.3 Statistics

All data processing and statistical analyses were performed in R (V4.0.5). Treatment and genotypic differences in growth responses in the BananaTainer were tested using the Kruskal-Wallis rank sum test, followed by pairwise Wilcoxon post-hoc tests (α = 0.05). Segmented regression on leaf area was performed to detect significant breakpoints in growth rate during the greenhouse experiments.

## 3 Results

### 3.1 Growth temperature responses are organ-specific

Ambient temperature affected plant organs in different ways, with shoot growth (leaves, pseudostem and rhizome) being more strongly influenced by a change in T regime than root growth. After 4 weeks at the 30 °C BananaTainer regimes, new leaves appeared significantly faster (p-value < 2.2 10^-16^), pseudostems were significantly taller (p-value < 2.2 10^-16^) and shoot fresh weight significantly increased (p-value = 2.5 10^-12^). Root fresh weight was not significantly affected by T regime, resulting in differentially distributed masses among the organs. While pseudostem:plant ratios were significantly increased (p-value < 2.2 10^-16^), root:shoot ratios significantly declined (p-value < 2.2 10^-16^) (Fig. 1).

**Fig. 1.**
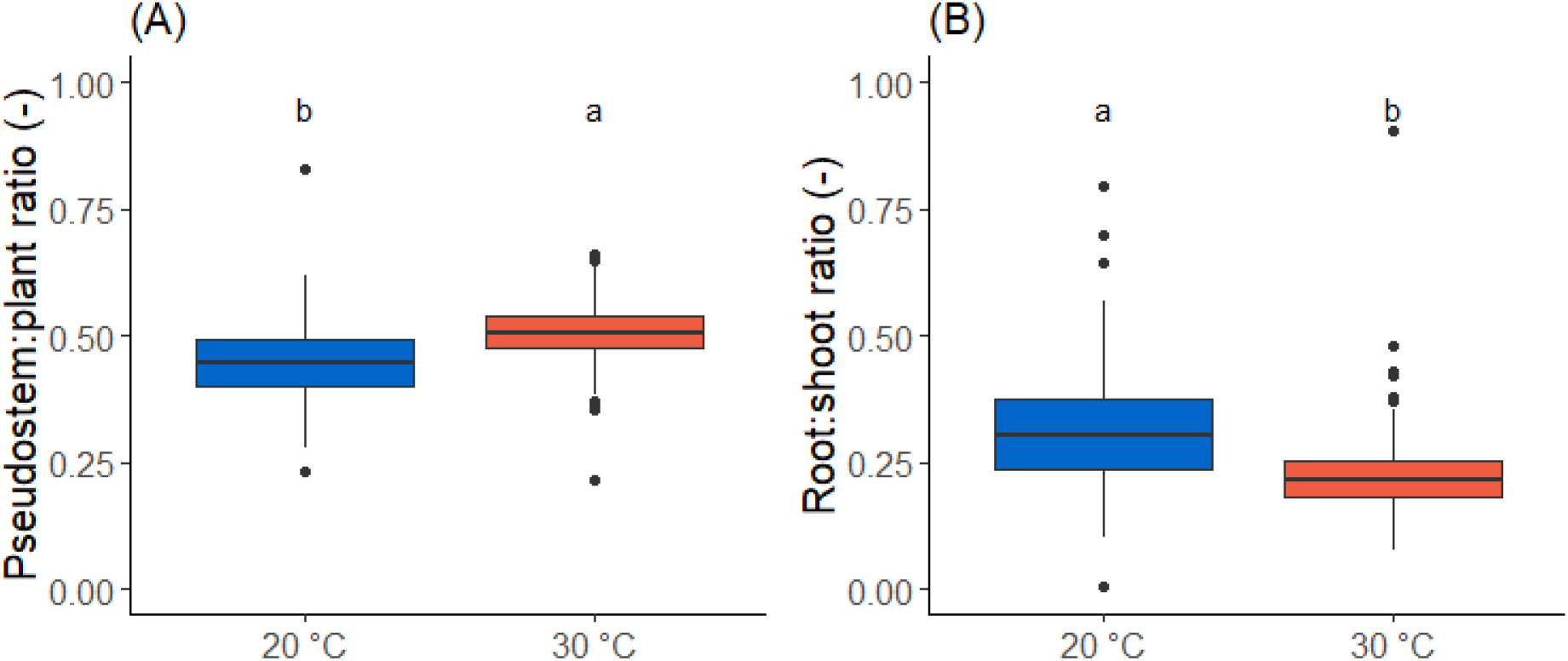
Organ-specific growth responses. Pseudostem:plant (A) and root:shoot ratios (B) at the end of the experiment of the 22 genotypes screened at 20 °C (blue) and 30 °C (red) in the BananaTainer. Different letters indicate significant differences (a > b; α = 0.05; n_20 °C_ = 642, n_30 °C_ = 614).

### 3.2 T_base_ is genotype-dependent, revealing diversity within subgroups

To determine the variability in T_base_, genotype-specific canopy growth models of the two reference genotypes were estimated. The average standard deviation of T_base_ was 1.4 °C. While Cachaco’s predicted T_base_ was 17.0 °C, it was extrapolated to 11.5 °C for Mbwazirume (Fig. 2).

**Fig. 2.**
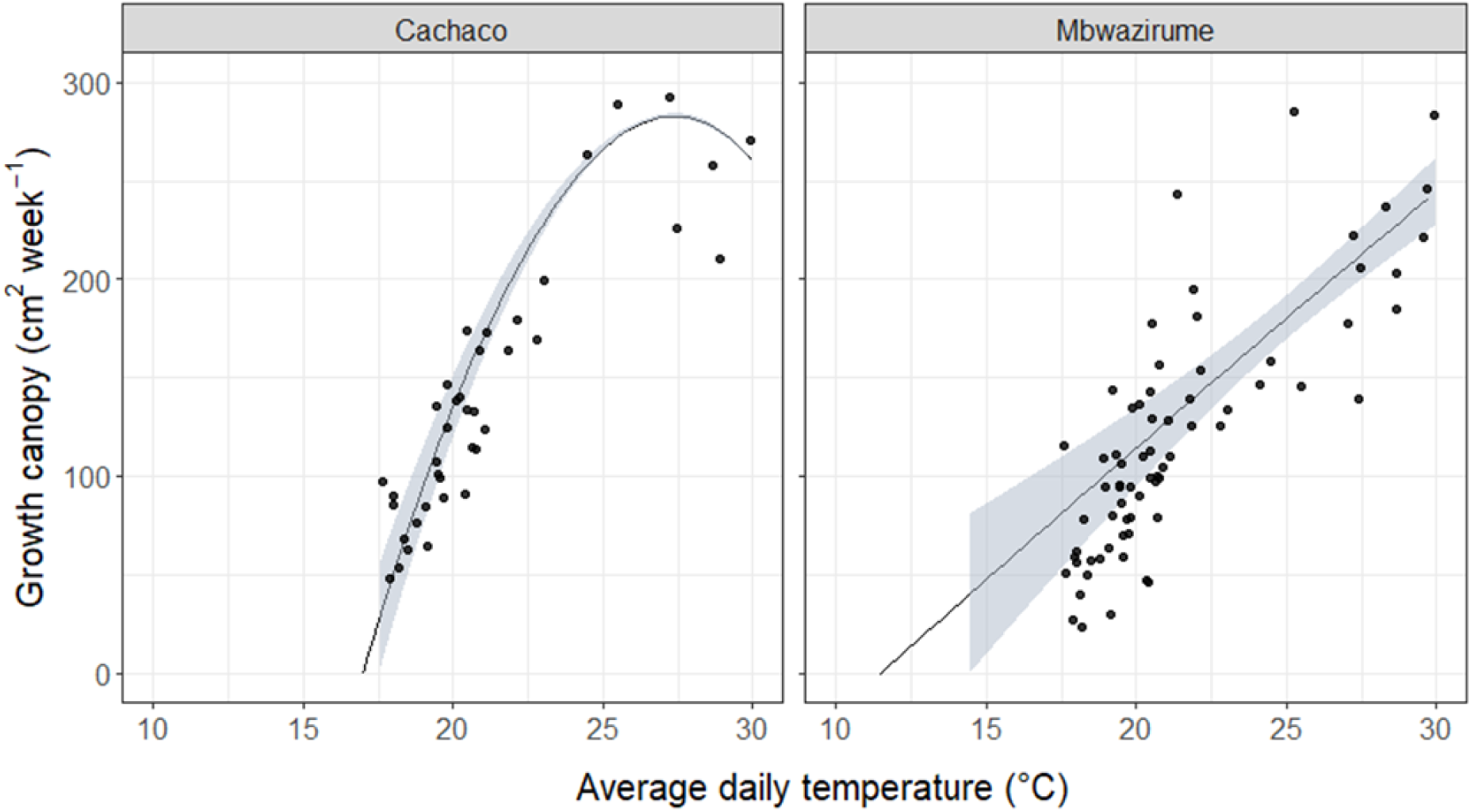
Modelled canopy growth curves of the reference genotypes (Eq. 2). Shaded area represents the standard deviation of the 5 and 12 growth models developed for Cachaco and Mbwazirume, respectively. Black dots show the average growth response per experienced temperature, based on a total of 411 Cachaco and 1058 Mbwazirume plants.

Among all screened genotypes, the average predicted T_base_ was 14.3 °C (± 4.1 °C). 35 of the 116 genotypes (30 %), had a predicted T_base_ below the generally accepted reference T_base_ of 14 °C (Fig. 3). T_base_ of the Mutika/Lujugira genotypes, endemic to the East African highlands, spanned a large T range. N’Dundu had one of the lowest predicted T_base_ (8.7 °C), while for Guineo T_base_ was predicted close to the reference (13.6 °C) (Fig. 3).

**Fig. 3.**
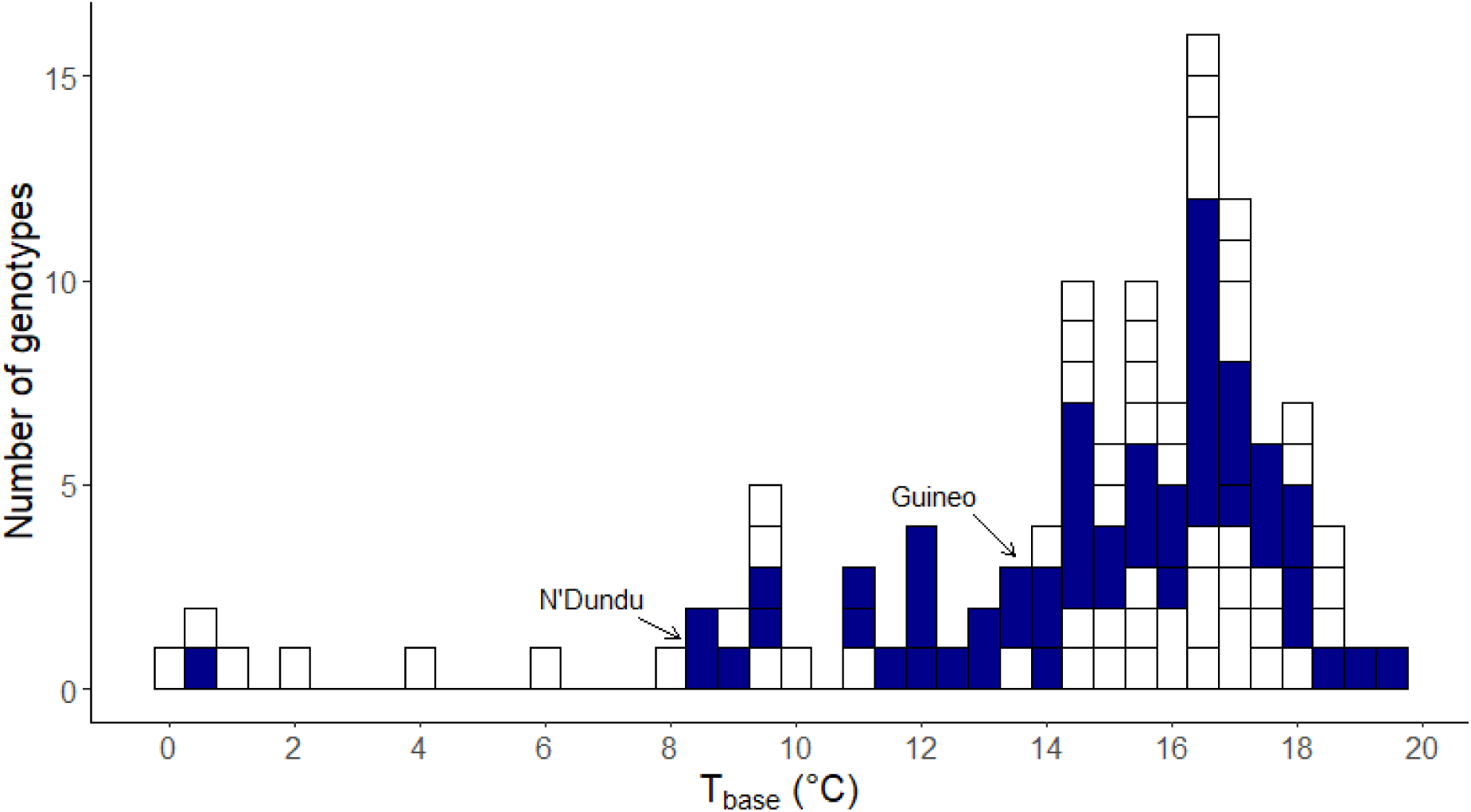
Histogram of the predicted base temperature (T_base_) of 116 genotypes. Dark blue bars represent the genotypes of the Mutika/Lujugira subgroup. Arrows refer to N’Dundu (T_base_ = 8.7 °C) and Guineo (T_base_ = 13.6 °C). Other rectangles correspond to different subgroups.

### 3.3 Greenhouse experiments validate the genotype dependent T_base_ and root:shoot ratios

During the greenhouse experiment with an average T of 17.1 °C, N’Dundu was the strongest growing genotype in terms of canopy, pseudostem and fresh plant growth (Fig. 4). N’Dundu showed an exponential growth response determined by its breakpoint in linear regressions 3 weeks after the start of the experiment (Supplementary Fig. S7). During the last 3 weeks, growth increased more strongly than during the first 3 weeks. Additionally, comparing with the 25.3°C greenhouse experiment, plants showed similar morphology changes, as the root:shoot ratio significantly decreased with increasing T (p-value = 2.3 10^-8^) (Supplementary Fig. S8). In this fluctuating environment both root and shoot fresh weight significantly increased with T (p-value = 6.3 10^-8^ and 3.3 10^-9^, respectively). On average, root fresh weight increased 2.5 times, while shoot fresh weight quadrupled.

**Fig. 4.**
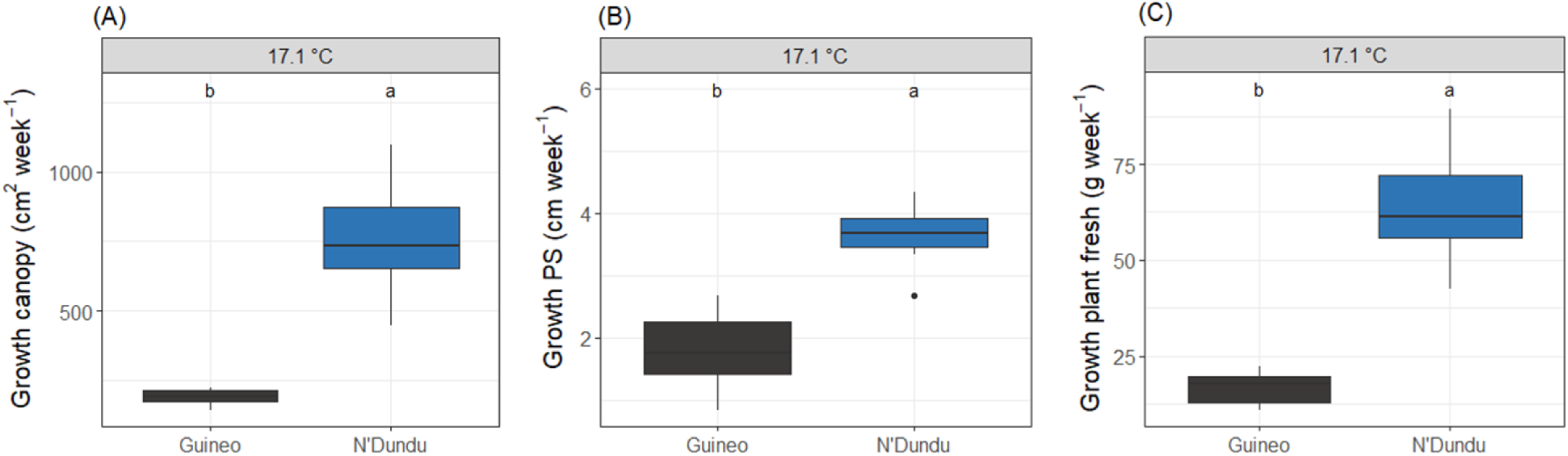
Growth response at the 17.1 °C greenhouse experiment. (A) Average weekly canopy, (B) pseudostem (PS) and (C) fresh plant growth at the 17.1 °C greenhouse experiment. Different letters represent significant genotypic differences (a > b; α = 0.05; n = 8).

### 3.4 The photosynthetic capacity plays a crucial role in temperature growth responses

Light- and CO_2_ response curves revealed significant increases in photosynthetic capacity (A_max_) and light saturation point (I_max_) with increased T (p-value < 10^-4^; data not shown), as a result of significant increases in the 3 rate limiting processes (V_cmax_, J_max_ and TPU_max_) (p-value < 10^-4^; data not shown). An increase in average T from 17.1 to 25.3 °C almost quadrupled V_cmax_ (data not shown), while J_max_ and TPU_max_ values almost tripled (data not shown).

Both genotypes responded significantly different at 17.1 °C (Fig. 5), with N’Dundu showing a significantly higher A_max_ and I_max_ (p-value = 0.04953) (Fig. 5B-C). Among the 3 rate limiting processes, V_cmax_, J_max_ and TPU_max_ were significantly higher in N’Dundu (p-value = 0.04953) (Fig. 5D-F). Additionally, transpiration efficiency (TE), defined as the increase in fresh plant weight per unit of water transpired, was significantly higher for N’Dundu at 17.1 °C (p-value = 0.00232) (Fig. 5A).

**Fig. 5.**
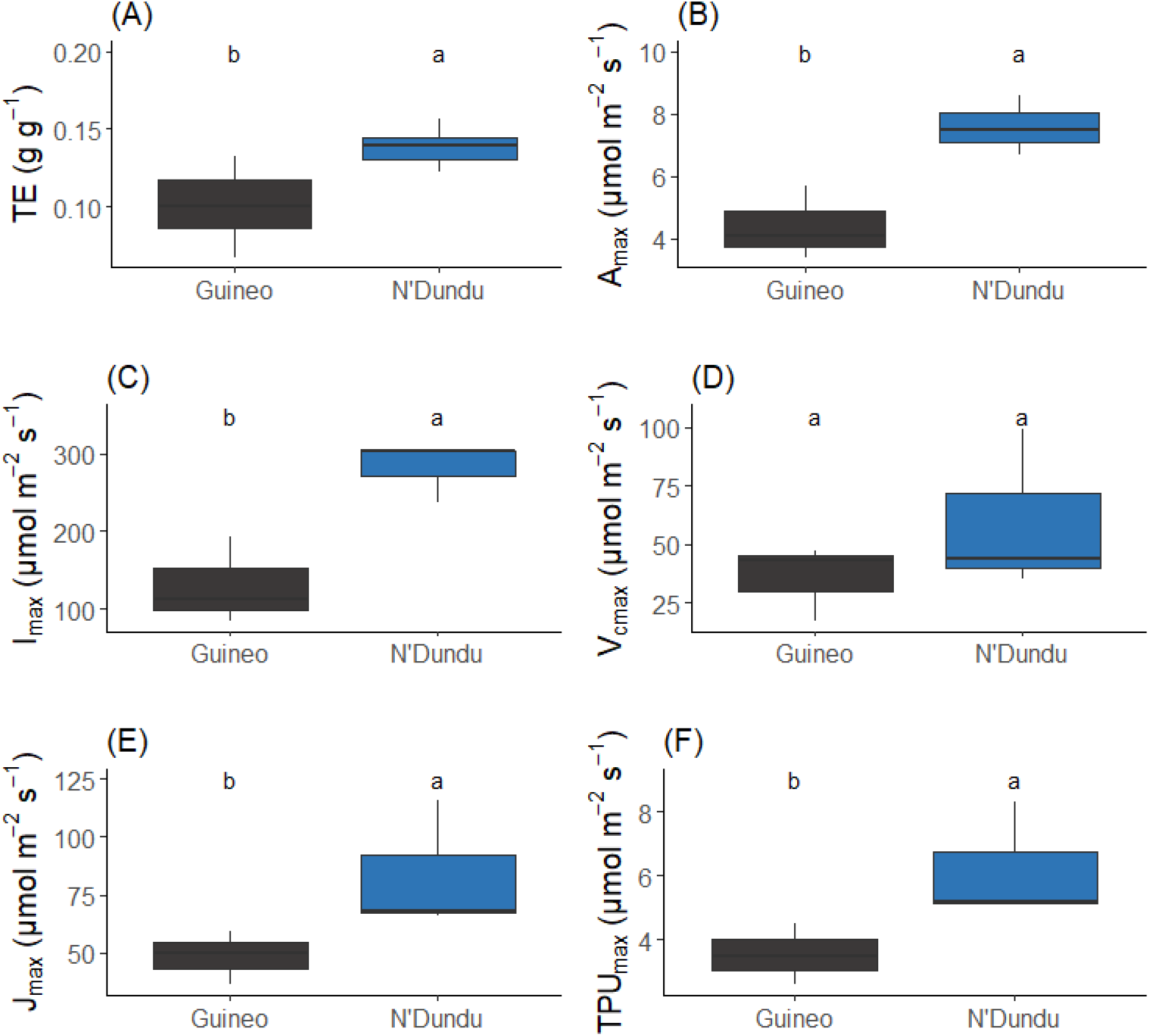
Genotype-specific biochemical capacity at 17.1 °C. (A) Transpiration efficiency (TE), (B) maximum photosynthetic rate (A_max_), (C) light saturation point (I_max_), (D) maximum carboxylation rate (V_cmax_), (E) maximum electron transfer rate (J_max_) and (F) maximum triose phosphate usage (TPU_max_) at average day T of 17.1 °C. Colours correspond to the genotype. Different letters indicate significant genotypic differences (a > b; α = 0.05; n = 3).

### 3.5 Future climates will boost plant development rates in the East African highlands

Despite the relatively low T_base_ of some genotypes, none of the screened genotypes are currently able to survive in the highest altitude regions of the East African highlands (Supplementary Fig. S9A). However, at lower located regions all screened genotypes would be able to grow as the average yearly temperature exceeds their genotype-specific T_base_ (Supplementary Fig. S9A). In Kawanda (Central Uganda), Guineo would currently accumulate 2900 GDD annually (Fig. 6A). N’Dundu would acquire 4700 GDD annually (Fig. 6E), accumulating as much GDD as Guineo in about 60 % of the time.

**Fig. 6.**
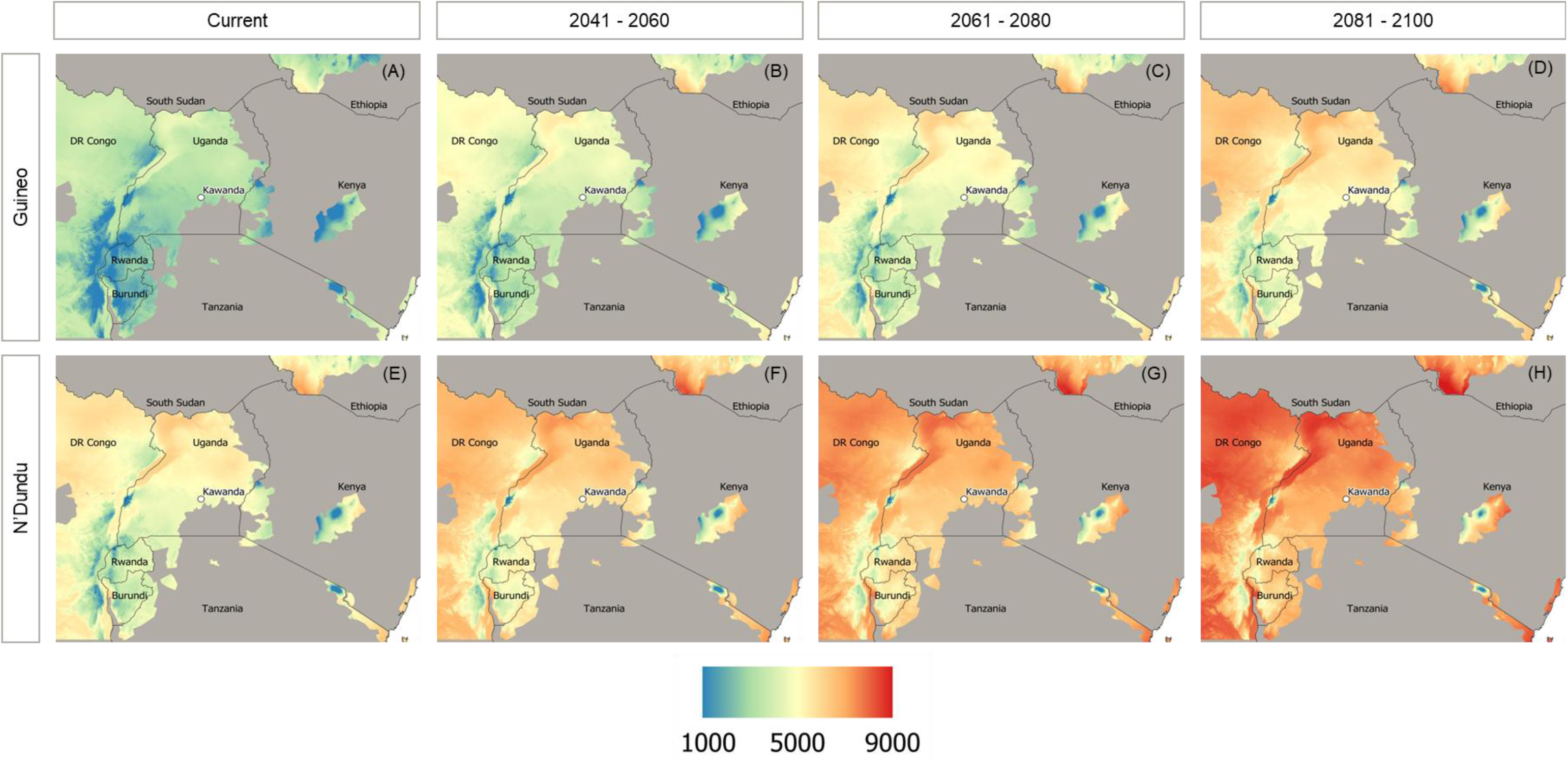
Current and future yearly GDD accumulation of Guineo and N’Dundu in the banana-growing region of the East African highlands. Upper row (A – D) represents the graphs for Guineo, the bottom row (E – H) for N’Dundu. From left to right, graphs represent the GDD accumulation in current climate (A, E), 2041 – 2060 (B, F), 2061 – 2080 (C, G) and 2081 – 2100 (D, H) (source: banana cultivating region from Ocimati et al. (2019); current and future climate from WorldClim (2022)).

Climate change, and its predicted increase in T, will also boost their development rates (Fig. 6B-D, F-H). According to the SSP5-8.5 emission scenario, mean annual T is expected to increase in Kawanda with 6.2 °C up to 27.5 °C by 2100. At this average daily T, Guineo and N’Dundu would accumulate 75 to 46 % more GDD as compared to the current environmental conditions, respectively (Fig. 6D,H). This way, they would acquire 5000 and 6860 GDD annually (Fig. 6D,H).

## 4 Discussion

### 4.1 Long-term exposure to stable temperature conditions reveals genotypic diversity in T_base_

The beta distribution model revealed with high precision genotypic diversity in terms of T_base_ (Fig. 2 and 3). Within species diversity in temperature growth responses is under debate (Parent and Tardieu, 2012). Turner and Hunt (1984) claimed that genotype-specific T_base_ exists and reported an estimated T_base_ range for 26 screened banana genotypes between 6 and 14 °C. However, the latter study used linear extrapolation to determine the genotype-specific T_base_. This method was later found to be less precise when used at T close to the optimal T (Yan and Hunt, 1999), thereby overestimating T_base_ during extrapolation. We confirm this genotype-specific T_base_ and reported that 20 of the 35 genotypes, that presented a T_base_ below the reference T_base_ of 14 °C, belonged to the most important subgroups on the African continent (Mlali, Mshare, Mutika/Lujugira and Plantain) (Supplementary Table S1). This considerable within subgroup variation in low T response was already observed by Turner et al. (2016) in plantains.

It is known that the Mutika/Lujugira genotypes have arisen by a cross between a *Musa acuminata* subsp. *banksii*/*Musa schizocarpa* hybrid and *Musa acuminata* subsp. *zebrina*, with a dominant contribution of *zebrina* (Christelová et al., 2017; Němečková et al., 2018). These hybridization events are thought to have occurred close to New Guinea and Java, from where the AA and AAA hybrids were introduced in East Africa during the first millennium (Perrier et al., 2019; Schoenbrun, 1993). Based on comparative linguistic evidence, it is assumed that banana reached the area of the Great Lakes region before the end of the first millennium (Perrier et al., 2019). Mitotic instability along with other somatic mutations, epigenetics and continuous selection by farmers have led to the current wide phenotypic diversity (Kitavi et al., 2020; Perrier et al., 2019; Shepherd and Da Silva, 1996; Šimoníková et al., 2020). This makes the East African highlands a secondary centre of origin, and the East African highland bananas are unique to the region.

### 4.2 Photosynthetic capacity contributes to the genotype-specific T suitability

Growth ranking of the 17.1 °C greenhouse experiment (Fig. 4) validated the predictions of superior growth for N’Dundu. Its photosynthetic capacity was about twice as high as that of Guineo, due to significant higher maximal triose phosphate usage (TPU_max_) and the capacity to process incident light by the photosynthetic apparatus J_max_ (Fig. 5). During photosynthesis, triose phosphates are exported from the chloroplast and converted to sucrose and starch, releasing inorganic phosphate which returns to the chloroplast to support further photosynthesis. TPU_max_ reflects the maximal triose phosphate use rate established by starch and sucrose synthesis and hence represents the capacity to regenerate inorganic phosphate. The TPU_max_ is known to be one of the major rate limiting photosynthetic processes at suboptimal T (Sage and Kubien, 2007). If sucrose and starch synthesis are too slow, metabolites accumulate and CO_2_ fixation is inhibited by a depletion of inorganic phosphate. Also J_max_ is of utmost importance to grow under suboptimal T. It reflects the capacity to regenerate ribulose 1,5- bisphosphate. As observed in many other tropical species, the combination of high light intensities and relatively low T more severely affects photosynthetic rates due to photoinhibition, resulting in rapid reduction of starch levels in sensitive species (Bongi et al., 1987; Taylor et al., 1972). This process of photoinhibition is expected to be of importance in subtropical and montane field conditions. Moreover, since it is very challenging to uncouple T and VPD and as nocturnal measurements neglect the impact of T on stomatal behaviour and consequently gas exchange and photosynthesis, it may explain why Parent and Tardieu (2012) were not able to show significant variability in T response within the same species.

### 4.3 Genotypic diversity in Mutika/Lujugira genotypes: a treasure for versatile performance in future tropical highlands

Temperature-wise, bananas are expected to thrive better in the East African highlands in future climates as compared to the current conditions (Fig. 6). Given that the average first crop cycle of banana varieties requires between 3,400 and 4,500 GDD (Lassoudiere, 2007; Turner, 1995), Guineo would be able to finish its crop cycle within 12 months by 2040 in Kawanda (Fig. 6D). In contrast, N’Dundu’s relatively low T_base_ would enable it to finish its crop cycle within a year under the current conditions if no other (a)biotic stress factors are at play (Fig. 6E). Important to note is that environmental conditions resulting in the most vigorous growth of a genotype are also considered most optimal for yield. In banana, biomass is strongly linked to bunch yield. Strong positive correlations between bunch weight and leaf emergence rate (Swennen and De Langhe, 1985), plant dry weight at harvest (Taulya et al., 2014), pseudostem volume at flowering (Stevens et al., 2020), pseudostem girth at flowering (Nyombi et al., 2009) and leaf area at flowering have been found (Aeberli et al., 2023).

Although the greenhouse experiment provides validation of two contrasting genotypes under more fluctuating conditions, validation in more agricultural relevant systems will be essential, especially since temperature changes are often intertwined with changes in other environmental parameters. Increased T in the highlands will coincide with increased evaporative demand (Muthoni et al., 2019). The majority of the Great Lakes Region currently has a bimodal rainfall with a short and a long dry season (van Asten et al 2011). Mutika/Lujugira genotypes have been proven to be ‘risk takers’, meaning they have a relatively low water use efficiency and keep a high transpiration rate for a longer period during the day (Eyland et al., 2021; Kissel et al., 2015; van Wesemael et al., 2019). Increased T will thus correspond to increased water demand. Calberto and colleagues estimated the increased water demand associated to climate change to range from 12 to 15 % (Calberto, Staver and Siles, 2015). Besides increased water demand, root:shoot ratios are predicted to decrease with increased mean annual T (Fig. 1B, Supplementary Fig. S8), creating an additional imbalance between water supply and demand. As already recommended by Manners et al. (2021), adapted farm management will have to be applied to fully exploit the benefits of climate change. Currently Mutika/Lujugira genotypes grow best at elevations between 900 and 1,200 m.a.s.l. (Kamira et al., 2016) and drought is already a major limiting factor in the area (van Asten et al 2011). Hence, cultivation of Mutika/Lujugira genotypes might need to shift to higher elevations (Supplementary Fig. S9), where evaporative demand is limited and excessive water loss avoided. Depending on the water management, farmers would need to opt for more drought tolerant genotypes, such as Simili Radjah and Cachaco (Eyland et al., 2021; van Wesemael et al., 2019). But those more hardy ABB types are not suitable to make matooke. Therefore breeding new East African highland bananas is also needed (Eyland et al., 2023).

## 5 Conclusion

This study reveals within species variation in T growth responses. Tropical highlands will benefit from the increased T associated with climate change. However, increased water demand in combination with decreased root:shoot ratios advocates for breeding more ‘conservative’ East African highland matooke types. To improve productivity of East African Highland Bananas, more attention should be given to research oriented towards improved water/drought stress management. High throughput phenotyping does not only provide useful information for gene banks, but also for breeding programs and will increase the implementation of suitable on-farm genotypic diversity. Screening of a wider range of genotypes will allow the selection of adapted genotypes to a given climate, thereby complying with the cultural and taste-specific requirements. We, therefore, recommend stakeholders to continue screening for T responses in both controlled and more agricultural relevant situations.

## Data availability statement

The raw data supporting the conclusions of this article will be made available by the authors, without undue reservation.

## Supplementary figures legends

**Fig. S1. The BananaTaine**r. A container based growth chamber with LED illumination. Banana plantlets are grown in trays per six, on three layers with a maximum capacity of 504 plants.

**Fig. S2.** Daily temperature regime for all temperature treatments on all BananaTainer levels. Blue colours correspond to a 20 °C run, green colours to a 25 °C run and red colours to a 30 °C run. Grey shaded area indicates dark period.

**Fig. S3.** (A) Original and (B) segmented topview picture of a banana plant to determine canopy leaf area. (C) and (D) represent the original and segmented picture of the individual leaves of the same banana plant, respectively. Red rectangle is the reference area of 50 cm^2^, the red arrow (7 cm^2^) indicates the youngest leaf at the start of the experiment.

**Fig. S4. Illustration of the beta distribution model.** T_base_, T_opt_ and T_max_ represent the base, optimal, and maximum temperature at which canopy growth occurs, respectively. Y_max_ is the maximum growth rate.

**Fig. S5. Decision tree to determine the accuracy of modelled growth curves.** Y_max_ represents the maximal growth rate, RMSE the root mean square error and RMSE_norm_ the normalized RMSE (RMSE/Y_max_). Models classified into the ‘YES’ category are considered satisfactory.

**Fig. S6.** Frequency histograms of the (A) temperature (T), (B) relative humidity (RH), (C) vapour pressure deficit (VPD) and (D) light intensity (PAR) observed during the 17.1 and 25.3 °C experiments between 08:00 and 20:00. Colours correspond to the T treatment.

**Fig. S7.** Leaf area time course along the different greenhouse experiments of Guineo and N’Dundu. Points represent the weekly average leaf area per genotype, error bars the standard deviation. Lines represent the best segmented fit for each genotype and temperature treatment. Colours correspond to the genotype (n = 8).

**Fig. S8.** Root:shoot ratio at the end of each greenhouse experiment after 6 weeks, validating the root:shoot ratio trend at increased temperature in the BananaTainer. Different letters indicate significant differences between temperature treatments (a > b; α = 0.05; n_17.1 °C_ = n_25.3 °C_ = 16). Colours correspond to the greenhouse temperature treatment.

**Fig. S9. Number of suitable genotypes in the banana growing region of the East African highlands.** Genotypes are considered suitable if T_base_ exceeds the average yearly T. (A) represents the number of genotypes in the current climate, (B) in 2041 - 2060, (C) in 2061 - 2080 and (D) in 2081 - 2100 (source: banana cultivating region from Ocimati *et al*. (2019); current and future climate from WorldClim (2022)).

## Acknowledgements

This high throughput phenotyping study was the result of a collaboration between the engineering team of Urban Crop Solutions and the laboratory of Tropical Crop Improvement at KU Leuven. The authors would, therefore, like to thank Jean-Pierre Coene, Maarten Vandecruys and Hendrik Siongers for their technical assistance to the hard- and software of the BananaTainer. The authors gratefully acknowledge Edwige Andre, Hendrik Siongers, Loïck Derette, Els Thiry, Hien Do, Barbara Grymonprez, Stan Blomme, Daan Van Gils, Simon Costers and Kaat Hebbelinck for the plant propagation and their assistance during plant growth and phenotyping.

CG and JvW were supported by a PhD scholarship funded by the Belgian Development Cooperation project “More fruit for food security: developing climate-smart bananas for the African Great Lakes region”. The authors thank all donors who supported this work through their contributions to the CGIAR Fund (http://www.cgiar.org/who-we-are/cgiar-fund/fund-donors-2/), and in particular to the CGIAR Research Program Roots, Tubers and Bananas (RTB-CRP) and to the *ERA-Net transnational call European Research Projects* LEAP Agri H2020 co-fund project on food & nutrition security & sustainable agriculture*, with funding from national funding agencies for the Project “PHENOTYPING THE BANANA BIODIVERSITY TO IDENTIFY CLIMATE SMART VARIETIES WITH OPTIMAL MARKET POTENTIAL IN AFRICA AND EUROPE”*

