## Supplementary figures for "Unlocking the power of gene banks: diversity in base growth temperature provides opportunities for climate-smart agriculture"

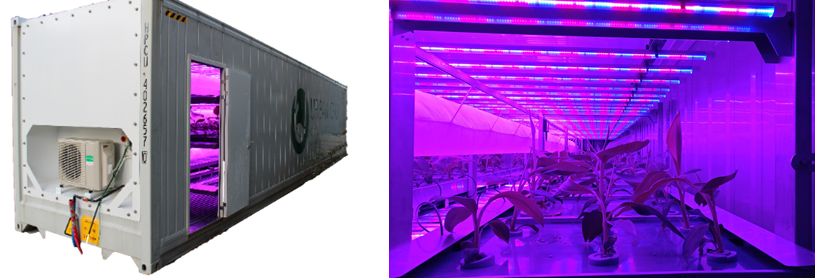
**Fig. S1. The BananaTaine**r. A container based growth chamber with LED illumination. Banana plantlets are grown in trays per six, on three layers with a maximum capacity of 504 plants.

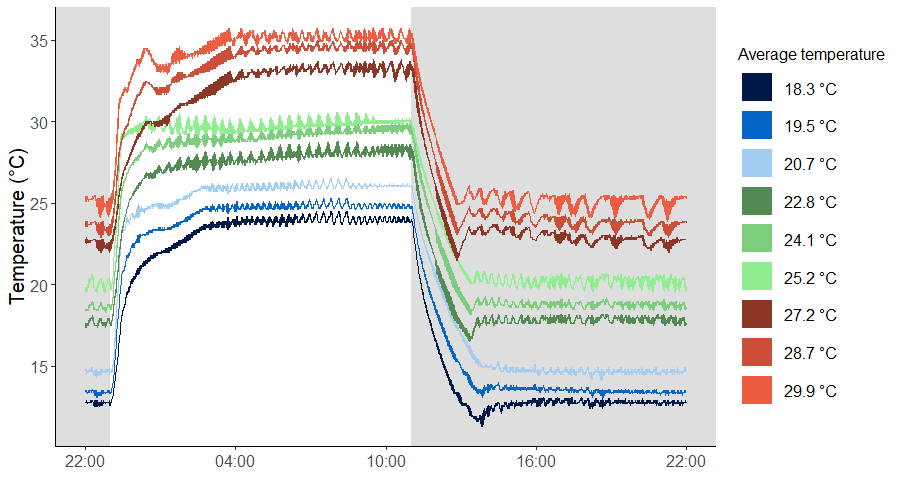
**Fig. S2.** Daily temperature regime for all temperature treatments on all BananaTainer levels. Blue colours correspond to a 20 °C run, green colours to a 25 °C run and red colours to a 30 °C run. Grey shaded area indicates dark period.

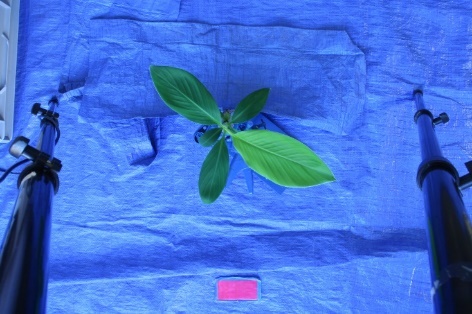

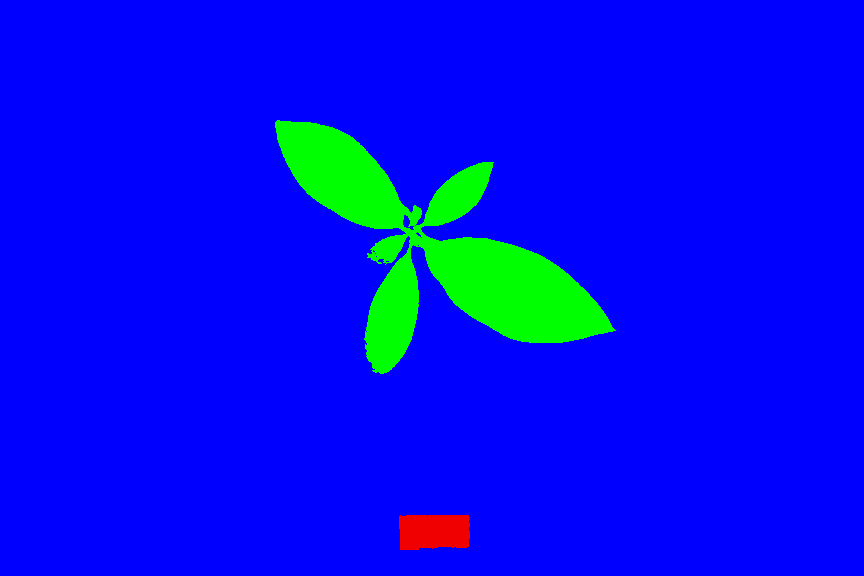

(A)

(B)

**Fig. S3.** (A) Original and (B) segmented topview picture of a banana plant to determine canopy leaf area.

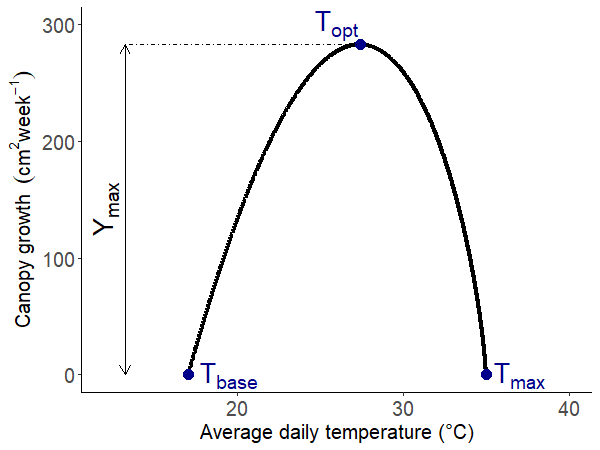
**Fig. S4. Illustration of the beta distribution model.** T_base_, T_opt_ and T_max_ represent the base, optimal, and maximum temperature at which canopy growth occurs, respectively. Y_max_ is the maximum growth rate.

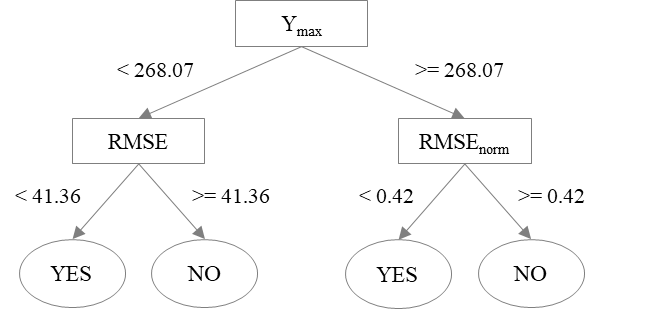
**Fig. S5.** **Decision tree to determine the accuracy of modelled growth curves.** Y_max_ represents the maximal growth rate, RMSE the root mean square error and RMSE_norm_ the normalized RMSE (RMSE/Y_max_). Models classified into the ‘YES’ category are considered satisfactory.

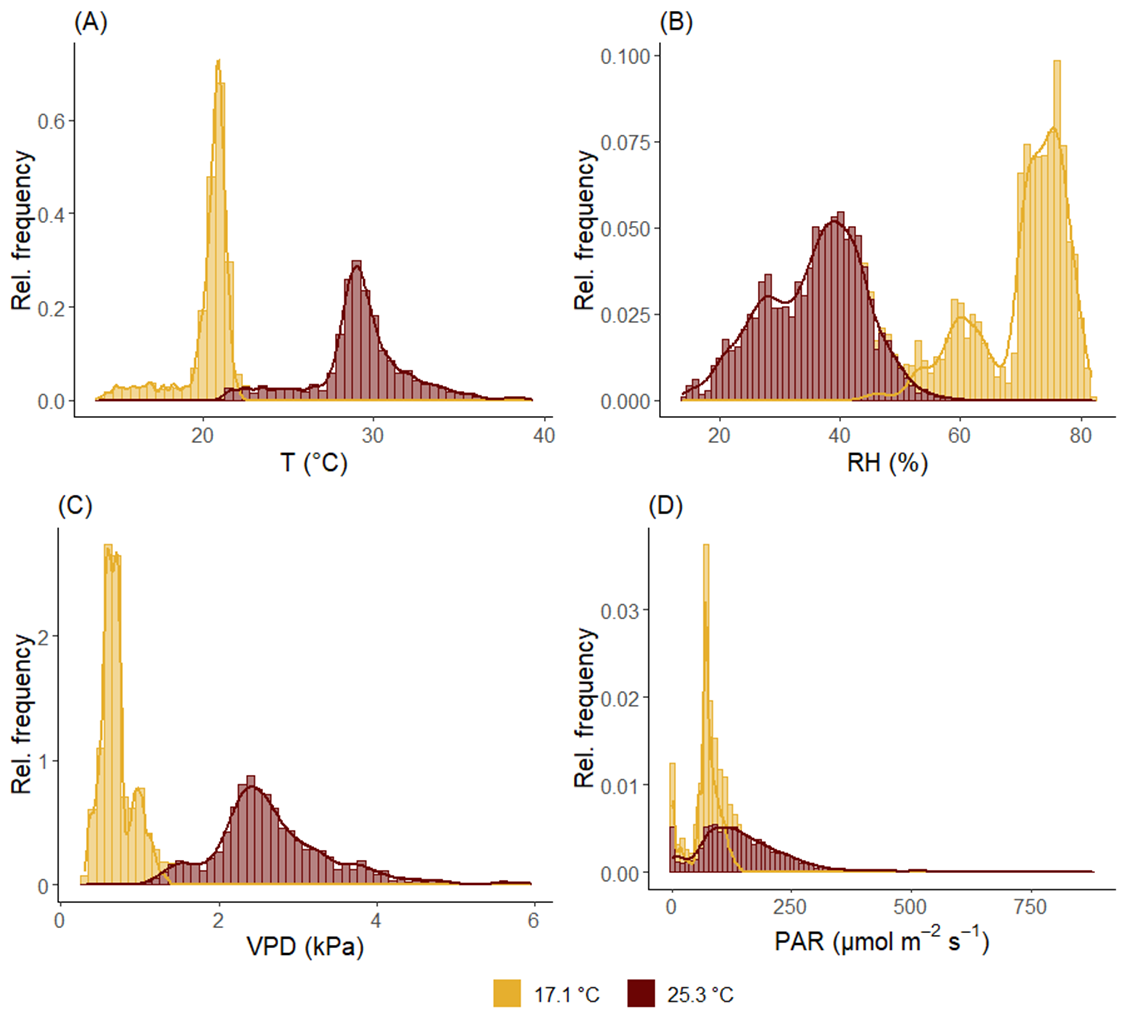
**Fig. S6.** Frequency histograms of the (A) temperature (T), (B) relative humidity (RH), (C) vapour pressure deficit (VPD) and (D) light intensity (PAR) observed during the 17.1 and 25.3 °C experiments between 08:00 and 20:00. Colours correspond to the T treatment.

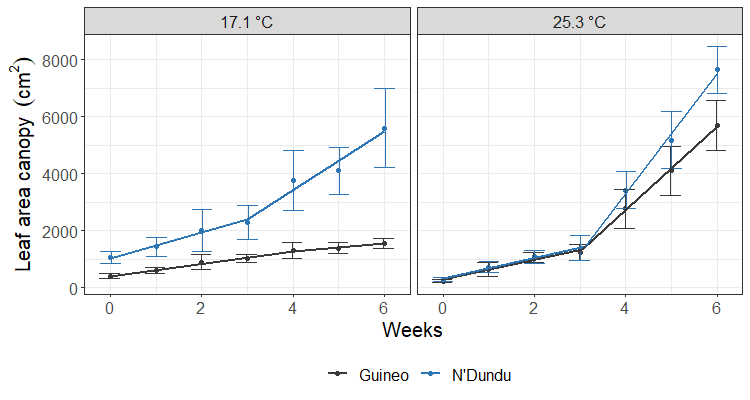
**Fig. S7.** Leaf area time course along the different greenhouse experiments of Guineo and N’Dundu. Points represent the weekly average leaf area per genotype, error bars the standard deviation. Lines represent the best segmented fit for each genotype and temperature treatment. Colours correspond to the genotype (n = 8).

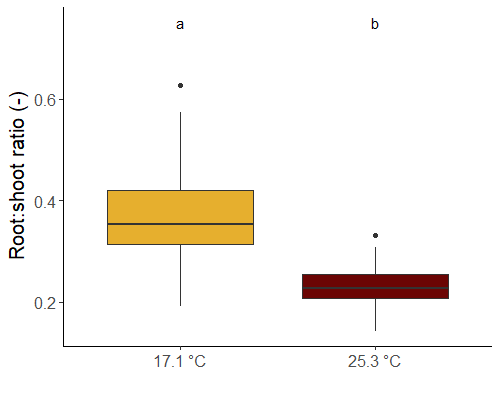
**Fig. S8.** Root:shoot ratio at the end of each greenhouse experiment after 6 weeks, validating the root:shoot ratio trend at increased temperature in the BananaTainer. Different letters indicate significant differences between temperature treatments (a > b; α = 0.05; n_17.1 °C_ = n_25.3 °C_ = 16). Colours correspond to the greenhouse temperature treatment.

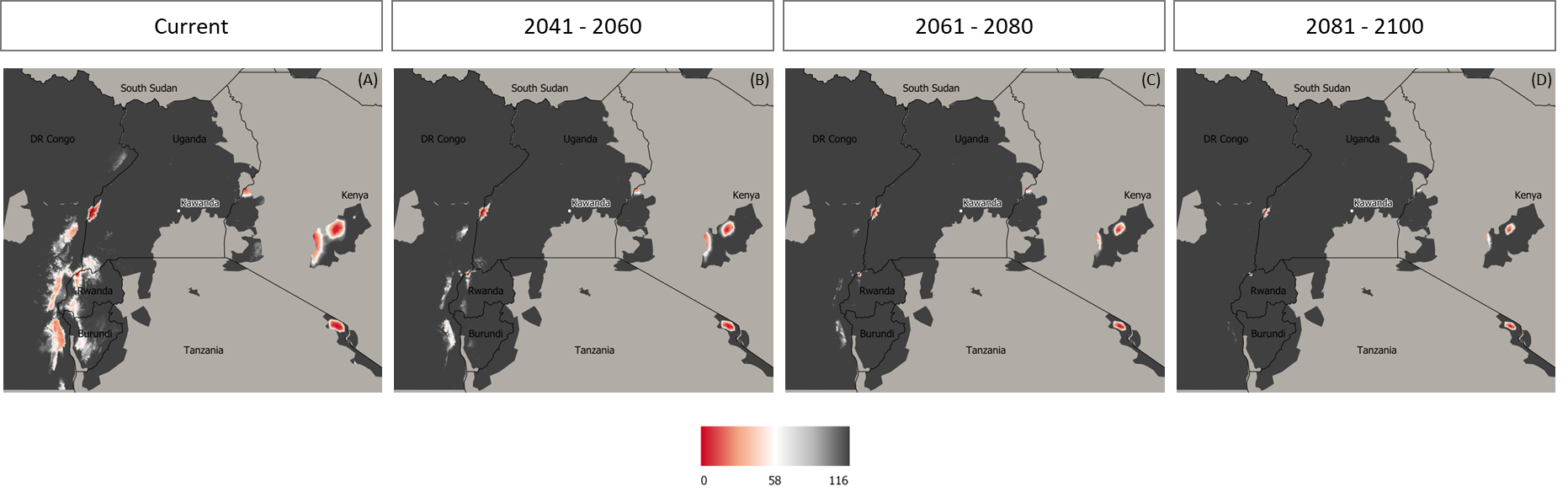
**Fig. S9.** **Number of suitable genotypes in the banana growing region of the East African highlands.** Genotypes are considered suitable if T_base_ exceeds the average yearly T. (A) represents the number of genotypes in the current climate, (B) in 2041 ‑ 2060, (C) in 2061 ‑ 2080 and (D) in 2081 ‑ 2100 (source: banana cultivating region from Ocimati *et al.* (2019); current and future climate from WorldClim (2022)).
